# A new framework for metabolic connectivity mapping using bolus [^18^F]FDG PET and kinetic modelling

**DOI:** 10.1101/2022.12.27.522050

**Authors:** Tommaso Volpi, Giulia Vallini, Erica Silvestri, Mattia De Francisci, Tony Durbin, Maurizio Corbetta, John J. Lee, Andrei G. Vlassenko, Manu S. Goyal, Alessandra Bertoldo

## Abstract

**Purpose:** Metabolic connectivity (MC) has been previously proposed as the covariation of static [^18^F]FDG PET images across participants, which we call *across-individual* MC (ai-MC). In few cases, MC has also been inferred from dynamic [^18^F]FDG signals, similarly to fMRI functional connectivity (FC), which we term *within-individual* MC (wi-MC). The validity and interpretability of both MC approaches is an important open issue.

Here we reassess this topic, aiming to 1) develop a novel methodology for wi-MC estimation; 2) compare ai-MC maps obtained using different [^18^F]FDG parameters (*K_1_*, i.e. tracer transport rate, *k_3_*, i.e. phosphorylation rate, *K_i_*, i.e. tracer uptake rate, and the standardized uptake value ratio, *SUVR*); 3) assess the interpretability of ai-MC and wi-MC in comparison to structural and functional connectivity (FC) measures.

**Methods:** We analyzed dynamic [^18^F]FDG data from 54 healthy adults using kinetic modelling to quantify the macro- and microparameters describing the tracer behavior (i.e. *K_i_*, *K_1_, k_3_*). We also calculated *SUVR*. From the across-individual correlation of *SUVR, K_i_, K_1_, k_3_*, we obtained four different ai-MC matrices. A new approach based on Euclidean distance was developed to calculate wi-MC from PET time-activity curves.

**Results:** We identified Euclidean similarity as the most appropriate metric to calculate wi-MC. ai-MC networks changed with different [^18^F]FDG parameters (*k_3_* MC vs. *SUVR* MC, r = 0.44). We found that wi-MC and ai-MC matrices are dissimilar (maximum r = 0.37), and that the match with FC is higher for wi-MC (Dice similarity: 0.47-0.63) than for ai-MC (0.24-0.39).

**Conclusion:** Our data demonstrate that individual-level MC from dynamic [^18^F]FDG data using Euclidean similarity is feasible and yields interpretable matrices that bear similarity to resting-state fMRI FC measures.

## Introduction

Positron emission tomography (PET) is one the earliest imaging modalities used for *in vivo* assessment of brain function, with very high specificity and increasingly high sensitivity [1]. In clinical practice, single-frame, static PET images are usually acquired, and the total amount of measured tracer activity is converted into a standardized uptake value ratio (*SUVR*). In experimental research settings, on the other hand, dynamic PET data are often obtained for full characterization of the tracer’s kinetic behavior through the lens of compartmental modelling [2].

In the neuroimaging literature of the recent decades, we have witnessed the rise of the field of connectomics, which aims at characterizing the structural and functional connections between brain areas using magnetic resonance imaging (MRI)-based techniques such as diffusion MRI (dMRI) and blood oxygen level-dependent (BOLD) functional MRI (fMRI) [3]. This field has opened new scenarios for brain PET as well, and in particular for the [^18^F]fluorodeoxyglucose ([^18^F]FDG) tracer, a glucose analog, which has been increasingly employed to estimate the so-called ‘metabolic connectivity’ (MC), describing the relationships between the metabolic states of different brain regions.

In most studies, only a group-level MC map is obtained. Typically, an atlas with predefined regions of interest (ROIs) is used to extract ROI-wise [^18^F]FDG semiquantitative values at individual level [4], and then their pairwise correlation across individuals is calculated (labeled here as ‘*across-individual* MC’ (ai-MC)). PET tracers other than [^18^F]FDG have also been employed, with reports typically talking of ‘molecular connectivity’ [5].

The ai-MC approach differs significantly, both in the calculation and in the interpretation, from what is typically done to estimate structural connectivity (SC) from dMRI, or functional connectivity (FC) from fMRI [3], where connectivity matrices are first derived at individual level, and then averaged across individuals to obtain group-level connectivity maps. Importantly, an individual estimate is necessary if one aims to use MC as a biomarker [6].

Only a handful of studies have attempted to use dynamic data to derive MC at individual level from PET time-activity curves (TACs), either in humans [6, 7, 8] or in animal models [9], an approach we label as *within-individual* MC (wi-MC). This is because handling PET *time series* comes with peculiar challenges as compared to fMRI time series or PET *subject series*, as we previously discussed [8]: the strong collinearity amongst TACs, which share the same positive trend related to [^18^F]FDG’s irreversible uptake, makes it impossible to directly employ simple correlation analysis as done with fMRI FC. To overcome this, previous works have opted for TAC standardization or detrending, not only in bolus injection protocols, but also for continuous infusion, where MC is becoming popular due to the higher temporal resolution of reconstructed infusion data [6, 9].

This approach, however, is problematic, especially with bolus injection studies, as it removes the main *signal* in PET TACs, and considers only the *fluctuations* around it, which may be related more to physical and statistical noise (measurement error [2]) than to biologically informative variability.

Thus, new solutions are highly warranted if wi-MC is to become useful and biologically interpretable.

Turning back to ai-MC approaches, we find that the information they have been based on so far is limiting, since all ai-MC studies use semiquantitative measures of [^18^F]FDG uptake (*SUVR*). Resorting to full kinetic modelling and a biologically validated compartmental description of [^18^F]FDG would provide important physiological information, such as the estimates of the tracer’s irreversible uptake (*K_i_*), which is correlated but not identical to *SUVR* [10], and the single rate constants, or microparameters, i.e., *K_1_* (tracer inflow, [ml/cm^3^/min]), *k_2_* (efflux [min^−1^]), *k_3_* (phosphorylation, [min^−1^]), which provide a self-consistent kinetic description of the initial steps of glucose metabolism [2, 11].

Using kinetic parameters instead of *SUVR* for ai-MC estimation may give rise to substantially different brain networks, but this endeavor has so far been hampered by the methodological difficulties associated with kinetic modelling, i.e., the need for long acquisitions and, ideally, arterial sampling to obtain an input function, the complexity of parameter estimation with noisy PET acquisitions etc. [2].

The validity and interpretability of MC measures has been the object of debate in the past. While validation of brain connectivity measures is a complex issue, a first simple readout can be provided by *between-modality association:* ai-MC maps, in particular, have been related both with SC [12] and with FC [13, 14]. From a biological standpoint, since SC represents the underlying axonal pathways between brain areas, we expect it to provide a “scaffold” for metabolic connections (as they should for FC [15]), and since FC is interpreted as representative of synaptic-level functional links between the activity of different neural populations [16], it also expected to be coupled with metabolic links, as glucose consumption happens mostly at the synapses [17].

Nonetheless, the evidence on the coupling between these network estimates is still sparse, and sometimes contradictory (for ai-MC vs. FC, see [13, 14]).

With these premises, we set out to provide a more comprehensive framework for [^18^F]FDG PET connectivity, using a large dataset of 54 healthy adults undergoing bolus dynamic PET studies.

First, to obtain wi-MC matrices, we compared different TAC standardization strategies and similarity metrics, and after choosing the best available approach, we proceeded to derive wi-MC matrices not only from *full* tissue TACs, but also from their *early* (0-10 min) and *late* part (40-60 min), to characterize MC networks more related to delivery (*early part*) or metabolism (*late part*) (**Figure 1**, *bottom*).

**Figure 1:**
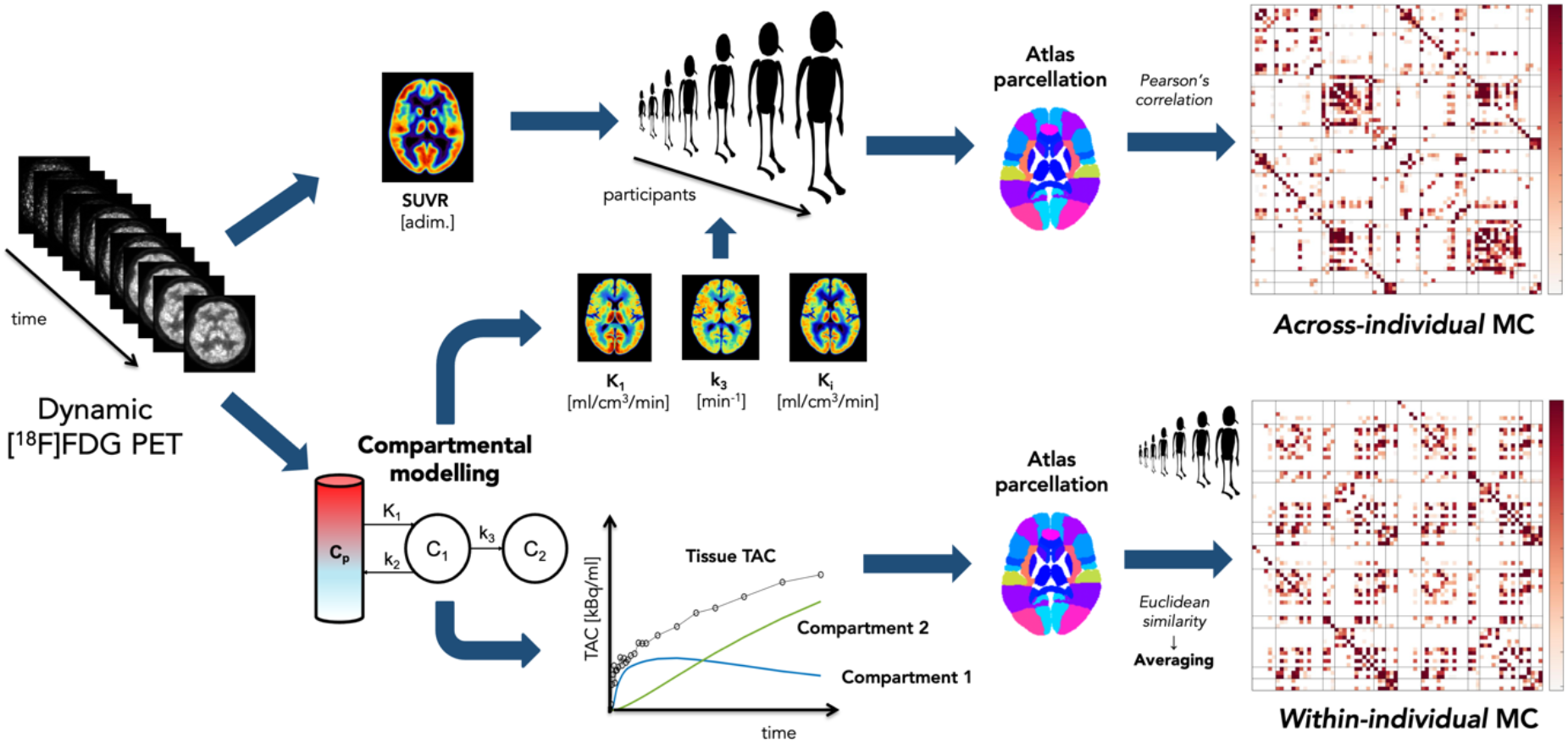
Analysis pipeline for inferring metabolic connectivity at the individual (*within-individual* approach) and group level (*across-individual* approach). [^18^F]FDG PET dynamic data (*far left*) is the source of all inferences of metabolic connectivity. A static *SUVR* image (*top left*) is obtained from frames in the 40-60 min window of the dynamic PET data; in parallel, compartmental modelling is fitted to dynamic PET data to estimate [^18^F]FDG kinetic parameters at the voxel level (using Variational Bayesian inference and an image-derived input function as a surrogate for the [^18^F]FDG plasma concentration *C_p_*), in particular *K_i_, K_1_* and *k_3_* (*center*), and reconstruct the time courses of compartments 1 and 2 (*bottom center*). From the *subject series* of *SUVR, K_i_, K_1_* and *k_3_*, parcellated thanks to the chosen atlas ROIs registered to individual PET space, we calculate across-individual MC via Pearson’s correlation (*top right*), while from the *time series* of the tissue TACs, compartments 1 and 2, individual-level MC is obtained via Euclidean similarity and averaged across participants (*bottom right*).

Then, we carried out [^18^F]FDG kinetic modelling, using Sokoloff’s two-tissue compartmental model [2, 11] and an image-derived input function (IDIF) calibrated with venous samples [18]. The ai-MC matrices were obtained from *SUVR, K_i_, K_1_*, and *k_3_*, describing the physiology of glucose kinetics (**Figure 1**, *top*). The reconstructed TACs of the first (*C*_1_(*t*), [kBq/cm^3^]) and second compartment (*C*_2_(*t*), [kBq/cm^3^]) of Sokoloff’s model, i.e., the time-varying intracellular concentrations of [^18^F]FDG and [^18^F]FDG-6P (representative of hexokinase activity), respectively, were used as additional inputs for wi-MC estimation.

Finally, to assess the validity and interpretability of wi-MC and ai-MC matrices, we compared them at multiple levels, i.e., in terms of similarity in matrix structure and ‘hub’ nodes (i.e., most connected nodes in each network [19]). The match of wi-MC and ai-MC estimates with other connectivity measures was also explored, i.e., SC from an average template [20] and FC from resting-state fMRI data of the same individuals.

## Materials and methods

### Participants

Fifty-four healthy adults (mean age 57.4 ± 14.8 years, 24 males) underwent [^18^F]FDG PET and MRI scans as part of the Adult Metabolism & Brain Resilience (AMBR) study [21]. Participants were excluded if they had contraindications to MRI, history of mental illness, possible pregnancy, or medication use that could interfere with brain function.

### Data acquisition

For each participant, structural and functional MRI acquisitions were performed on a Siemens Magnetom Prisma^fit^ scanner, while [^18^F]FDG scans were performed on a Siemens model 962 ECAT EXACT HR + PET scanner (Siemens/CTI), with concomitant venous blood sampling. For details on these acquisitions, see **Supplementary Methods**.

### MRI preprocessing

Structural T1w images were preprocessed as reported in **Supplementary Methods**.

The Hammers anatomical atlas [22] and the Schaefer functional atlas (100 ROIs, 7 networks) [23] were registered to T1w.

For the *Hammers* atlas, 74 ROIs (out of 83) were kept for further analysis, after removing white matter (WM) and cerebrospinal fluid (CSF)-only ROIs. For simpler visualization and interpretation, the regions were divided into 7 anatomical clusters, i.e., 1) frontal lobe, 2) temporal lobe, 3) parietal lobe, 4) occipital lobe, 5) insula and cingulate gyri, 6) subcortical structures, 7) cerebellum. The complete list of the regions (in the same order as in the manuscript’s figures) is reported in the **Supplementary Methods**.

For the *Schaefer* atlas, the 100 ROIs were supplemented by 12 subcortical ROIs taken from the Hammers atlas (bilateral caudate, accumbens, putamen, pallidum, thalamus, cerebellum).

The rs-fMRI data were preprocessed as reported in the **Supplementary Methods**.

Pre-processed rs-fMRI signals were obtained within each ROIs from the Hammers and Schaefer atlases, which had been linearly mapped from T1w to rs-fMRI space, by averaging over voxels within the T1w grey matter (GM) segmentation (probability > 0.8 of belonging to GM).

### PET kinetic modelling

Dynamic PET data were motion-corrected using FSL’s *mcflirt* [24].

A static PET image was obtained by summing motion-corrected late PET frames (40-60 min). The static image was linearly registered to T1w space using FSL’s *flirt* [24], and normalized into *SUVR* dividing by the whole-brain [^18^F]FDG average uptake [25].

To perform kinetic modelling, an IDIF approach was used, and voxel-wise estimation of Sokoloff’s model parameters was performed using a Variational Bayesian approach [26]. Parametric maps (i.e., voxel-wise maps) of *K_1_, k_2_, k_3_, V_b_* [%] (blood volume fraction), and *K_i_* were obtained for each participant. The voxel-wise predicted time courses of *C*_1_(*t*) and *C*_2_(*t*) were reconstructed from Laplace transform solutions of Sokoloff’s model [11] (see **Supplementary Methods**).

The group-average maps of *SUVR, K_i_, K_1_, k_3_* are shown in **Supplementary Figure 6**.

### Within-individual metabolic connectivity (wi-MC)

ROI-level PET signals ([^18^F]FDG tissue TACs, *C*_1_(*t*) and *C*_2_(*t*)) were extracted from the Hammers and Schaefer regions, which had been linearly mapped from T1w to PET space, by averaging over voxels within the GM segmentation (probability > 0.8 of belonging to GM).

The GM segmentation, being quite conservative, produces an average sample TAC which minimizes partial volume effects (PVEs) [27]. Moreover, spatial smoothing of the PET data during processing was avoided, further minimizing PVEs, as also suggested in many recent PET analysis reports [28].

The five-second frames of the ROI-wise tissue TACs (120 s in total) were temporally filtered by averaging them in triplets. Denoising was not performed on the TACs of *C*_1_(*t*) and *C*_2_(*t*), which are noise-free since they are derived from compartmental modelling. All the signals (tissue TACs, *C*_1_(*t*) and *C*_2_(*t*)) were then interpolated on a uniform virtual grid (five-second step).

To calculate wi-MC, several methods for standardizing PET TACs and evaluating their relationship (Pearson’s correlation, Cosine similarity, Euclidean similarity) were tested and compared (**Supplementary Figure 1, Supplementary Methods**) on the basis of a) their capability of retrieving a ‘biological’ network structure having subnetworks along the main diagonal, and inter-hemispheric connections between homologous regions [19], b) the between-individual variability of the edges of the MC network.

wi-MC matrices were calculated from:

a. the *full* tissue TACs (0-60 min, all 52 data points),
b. the *early* part of the tissue TAC (0-10 min, first 38 data points),
c. the *late* part of the tissue TAC (40-60 min, last 4 data points),
d. the full TACs of *C*_1_(*t*),
e. the full TACs of *C*_2_(*t*),

and then averaged across individuals into five group-level wi-MC matrices.

The between-individual variability of each wi-MC approach was calculated at edge level as the coefficient of variation (CV%), i.e., the percent ratio of the median absolute deviation (MAD) of the Euclidean similarity (ES) values across individuals divided by the median. An overall index of the between-individual variability was obtained by taking the median ± MAD of the CVs% for each matrix (**Supplementary Figure 5)**.

The association between each pair of wi-MC matrices was estimated using Pearson’s correlation, calculated between the upper triangular portions of each matrix, both without sparsification and after imposing a threshold (80^th^ percentile sparsity), as advocated in connectivity studies [30]. The sparse results are presented in the paper (full matrix results are in the **Supplementary Materials**). The significance of the correlation coefficients was assessed using Mantel’s test [31], which was tested for significance by 15,000 permutations. Additionally, p-values were Bonferroni corrected for 10 comparisons.

### Across-individual metabolic connectivity (ai-MC)

The [^18^F]FDG parametric maps were parceled at individual level with the Hammers and Schaefer atlas. The region-wise *SUVR, K_i_, K_1_, k_3_* values were within-individual normalized via z-scoring, in accordance with [4]. ai-MC matrices for *SUVR, K_i_, K_1_, k_3_* were computed with Pearson’s correlation (**Figure 1**, *top*).

The association between each pair of ai-MC matrices was estimated using Pearson’s correlation (upper triangular, with and without sparsification, with Mantel significance testing, and 6-fold Bonferroni correction).

### Comparing wi-MC vs. ai-MC: matrices, hubs and graph metrics

The information provided by the *wi-MC* vs. *ai-MC* approaches was compared via multiple strategies: at *edge* level by calculating Pearson’s correlation between the matrix elements, and at *region* level by comparing the hub nodes of each MC matrix.

For direct matrix-to-matrix comparison, Pearson’s correlation coefficients were calculated (upper triangular portions of MC matrices, with and without sparsification, with Mantel significance testing, and 20-fold Bonferroni corrections).

To identify hub nodes, all matrices were thresholded at the 80^th^ percentile, as previously advocated [30]. Then, hubs were identified on each matrix as the nodes belonging to the top 20% of the joint graph theory metrics degree (DEG) and eigenvector centrality (EC), thereby identifying nodes with high local and global connectivity [19].

For comparison of hubs across matrices, the Dice similarity between pairs of binary hub/non-hub vectors of wi-MC and ai-MC was assessed.

In addition, the regional EC values from both wi-MC and ai-MC matrices were plotted against [^18^F]FDG parameters (*SUVR, K_i_, K_1_, k_3_*) averaged across individuals (**Supplementary Results**).

### Matching wi-MC and ai-MC with structural and functional connectivity

The SC template was derived using 178 tractographies made available in the BCBToolkit as Disconnectome 7T [32]. Individual tractographies were reconstructed using the HCP 7T diffusion-weighted imaging dataset [20]. Single-individual SC matrices were calculated using DSI studio (https://dsi-studio.labsolver.org/). The final template was derived by averaging the connectivity matrices across controls. For consistency with MC, the sparsity level of the matrix was set to 80%. The MC vs. SC analysis was reproduced on a different publicly available tractography atlas [33] (**Supplementary Figure 12**).

For each participant, the functional connectivity (FC) matrix was obtained by means of Pearson’s correlations computed between fMRI time series of each pair of ROIs. Motion-corrupted volumes having frame-wise displacement higher than 0.4 mm were censored before FC computation [34]. FC matrices were then Fisher z-transformed and averaged across individuals to obtain the group-level FC. As with MC, the sparsity level was set at 80%.

To assess the agreement between the estimated metabolic connections and the SC and FC structure, the Dice similarity between binarized SC and each binarized wi-MC (group average) and ai-MC network was computed.

## Results

### Within-individual MC maps from PET time courses

We started by comparing different MC estimation methods (Euclidean similarity, Pearson’s correlation, Cosine Similarity) and TAC standardization approaches (**Supplementary Figure 1**), and verified that the Euclidean similarity method was the only one capable of retrieving structured MC matrices even without any signal normalization: in particular, in all the matrices reported in **Figure 2** (group-average wi-MC for the *Hammers* atlas, with 80% sparsity; **Supplementary Figure 2**, without sparsity-inducing thresholds; **Supplementary Figure 3** for the *Schaefer* functional atlas), we clearly see both a) a block-diagonal structure along the main diagonal, and b) enhanced secondary diagonals, representing intra-hemispheric within-‘network’ connections and inter-hemispheric homotopic connections (i.e., between homologous regions) respectively. Therefore, we exclusively present wi-MC matrices as obtained by the Euclidean similarity approach.

**Figure 2:**
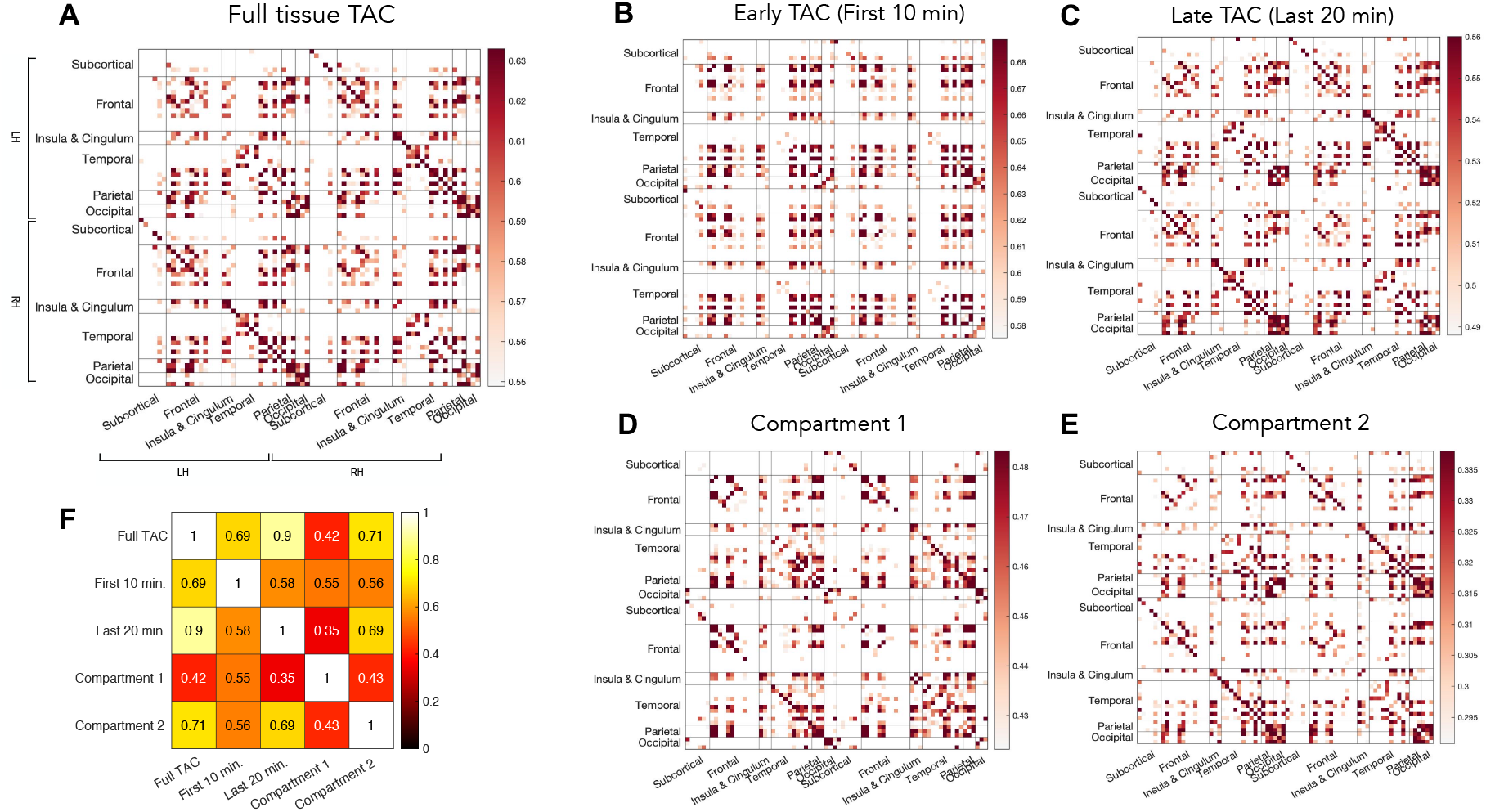
Matrix representation of *within-individual* metabolic connectivity. Matrix elements indicate Euclidean similarity of tracer kinetics from dynamic PET data of different regions. The Hammers parcellation is used to sample dynamic PET time series at the individual level. Following matrix computation for individual subjects, we averaged matrices for the group of 54 individuals. The average matrices for the *full* tissue TAC (**A**), its *early* part (**B**) and *late* part (**C**), the kinetics of *C*_1_(*t*) (**D**) and *C*_2_ (*t*) (**E**) are displayed. For clarity and for conforming to conventions for identifying network hubs, matrices are visualized with 80% enforced sparsity. Homotopic inter-hemispheric connections are most evident in (**A**) and (**C**), i.e., matrices which are more related to *metabolic* events, while in (**B**), which highlights the early, *flow*-related information, there is little difference between inter-hemispheric and intra-hemispheric connections. Pearson’s correlation coefficients between the upper triangular elements of the five wi-MC matrices are also reported (**F**), confirming the full tissue TACs associate most with the late part.

When visually evaluating the wi-MC matrices from the *full* TAC (**Figure 2A**), the *late* part (**Figure 2C**), and *C*_2_(*t*) (**Figure 2E**), areas of strong within-‘network’ relationships are located in the frontal and occipital cortex, but also in medial temporal lobe.

However, this network structure is clearly modified in the wi-MC from the *early* part of the TAC (**Figure 2B**), and *C*_1_(*t*) (**Figure 2D**): the occipital lobe loses ‘connectivity’, and the temporal and parietal areas become highly connected both within and between ‘network’. Moreover, the early TAC matrix seems to show a marked decrease of homotopy.

Subcortical structures tend to always display lower wi-MC than cortical areas (see **Supplementary Figure 4**).

The between-individual variability of the obtained wi-MC matrices (**Supplementary Figure 5**) is overall low, being lowest for the *full* TAC (CVs% median ± MAD of the entries of the upper triangular portion: 8.3% ± 28.9%), and highest for *C*_2_(*t*) (CVs% median ± MAD of the entries of the upper triangular portion: 47.7% ± 9.4%).

The Pearson’s correlation between the average wi-MC matrices shows that the *full* TAC wi-MC has strong correlations with the *early* part, *C*_2_(*t*) and especially the *late* part of the TAC, as it is expected: the tails of the TACs have higher amplitudes, and thus have the strongest contribution to the between-region similarity in Euclidean sense. Notably, the *C*_1_(*t*) MC has instead weak relationships with the other wi-MC matrices, except for a moderate correlation (r = 0.55, Mantel’s test, p < 10^−9^, Bonferroni corrected) with the *early* TAC (**Figure 2F**).

In summary, by using PET TACs and the Euclidean similarity metric, it is possible to obtain individual-level MC matrices showing both within-‘network’ and homotopic connections, characterized by a low between-individual variability, and with the additional possibility of highlighting different physiological events (by considering the *full* kinetics, a *portion* of it, or the kinetics of the model compartments). Importantly, these features are present for both Hammers and Schaefer atlas.

### Across-individual MC maps: SUVR and kinetic parameters

The ai-MC matrices are displayed in **Figure 3** (full matrix version in **Supplementary Figure 7**; Schaefer version in **Supplementary Figure 8**).

**Figure 3:**
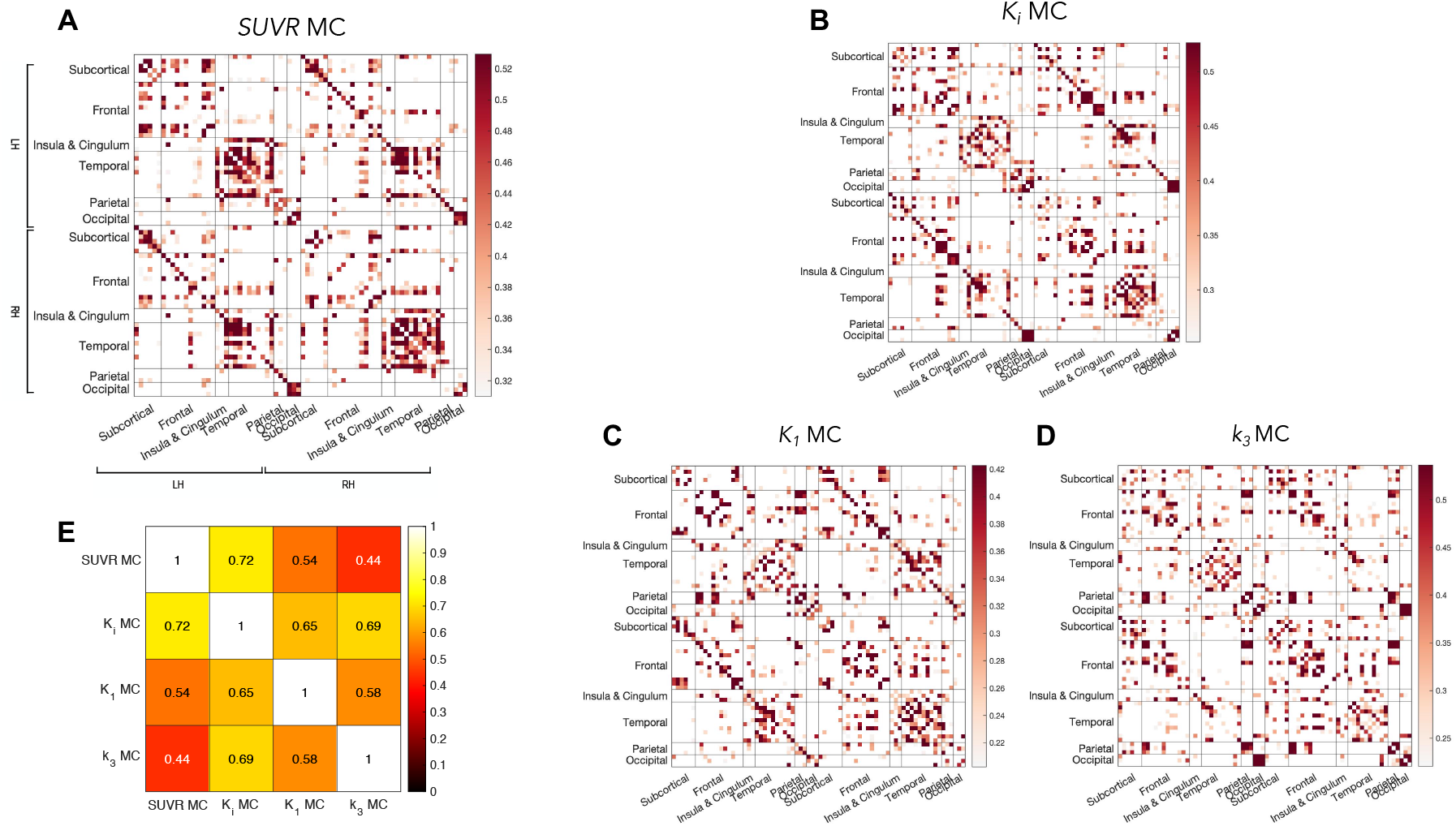
Matrix representation of *across-individual* metabolic connectivity. Matrix elements indicate Pearson’s correlation coefficients between [^18^F]FDG parameter values of different regions across our group of 54 individuals. The Hammers parcellation is used to sample individual-level [^18^F]FDG parametric maps. The across-individual MC matrices for *SUVR* (**A**), *K_i_* (**B**), *K_1_* (**C**) and *k_3_* (**D**) are reported. For clarity and for conforming to conventions for identifying network hubs, matrices are visualized with 80% enforced sparsity. The Pearson’s correlation coefficients between the upper triangular elements of the four ai-MC matrices are also reported (**E**), highlighting how *k_3_* MC has the lower similarity to traditionally calculated *SUVR* MC.

The ai-MC of [^18^F]FDG kinetic model parameters (*K_i_*, *K_1_, k_3_*) is presented here for the first time (see also *k_2_* and *V_b_* ai-MC matrices in **Supplementary Figure 9**) while ai-MC based on *SUVR* is the most frequently used approach in literature.

Some similarities are shared between the different parameters, especially between MC of *SUVR* and *K_i_* (r = 0.72, Mantel’s test, p < 10^−9^, Bonferroni corrected), with strong within-‘network’ connections in temporo-limbic areas. The *k_3_*-based ai-MC is instead quite different from the others, with enhanced connections in frontal areas and subcortical structures, and is in fact the least correlated with the others, especially with *SUVR* MC (r = 0.44, Mantel’s test, p < 10^−9^, Bonferroni corrected).

### Comparing within-individual vs. across-individual MC: similarity of matrices and hubs, match with [^18^F]FDG parameters

When the wi-MC and ai-MC networks were related via Pearson’s correlation (**Table 1**), some significant relationships are found, especially between wi-MC matrices and *K_1_* and *k_3_* ai-MC. However, the correlation values are generally low, with a maximum r = 0.37; even lower values are found for the Schaefer atlas (maximum r = 0.28). *SUVR* ai-MC seems to carry no meaningful relationships with wi-MC approaches.

**Table 1:**
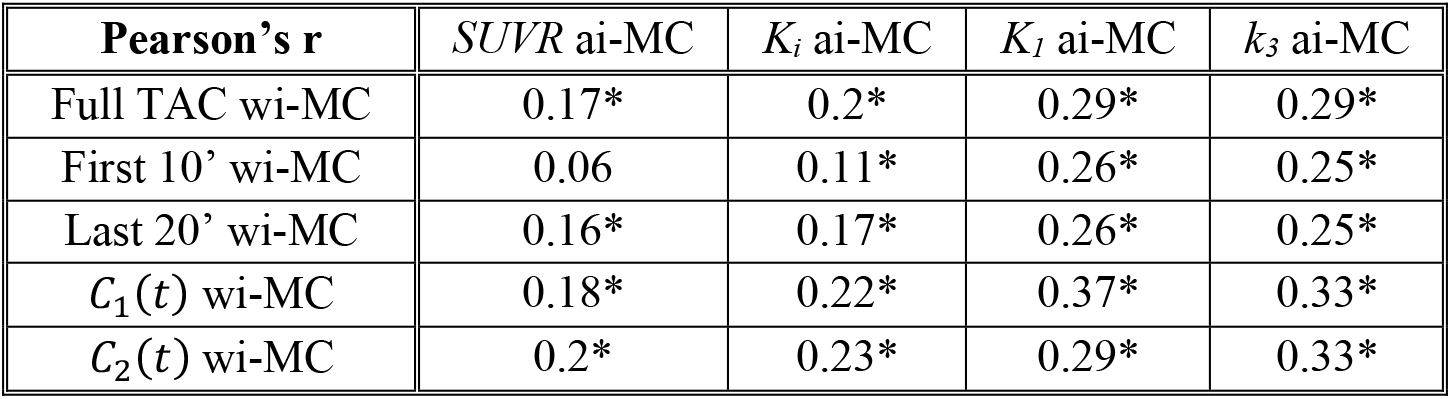
Within-individual vs. across-individual MC correlations. Across-edge Pearson’s correlations between *within-individual* (group-average, rows) and *across-individual* (columns) MC matrices (upper triangle, 80^th^ percentile threshold). Significant correlations (Mantel’s test, p < 0.05, Bonferroni corrected) are reported as *.

We then identified ‘hub’ nodes, i.e., highly connected nodes in each network, as is typically done in the field of connectomics.

wi-MC hubs are mainly located in frontal and temporal areas, with the exception of *C*_1_ (*t*) with more parietal involvement (**Figure 4A**).

**Figure 4:**
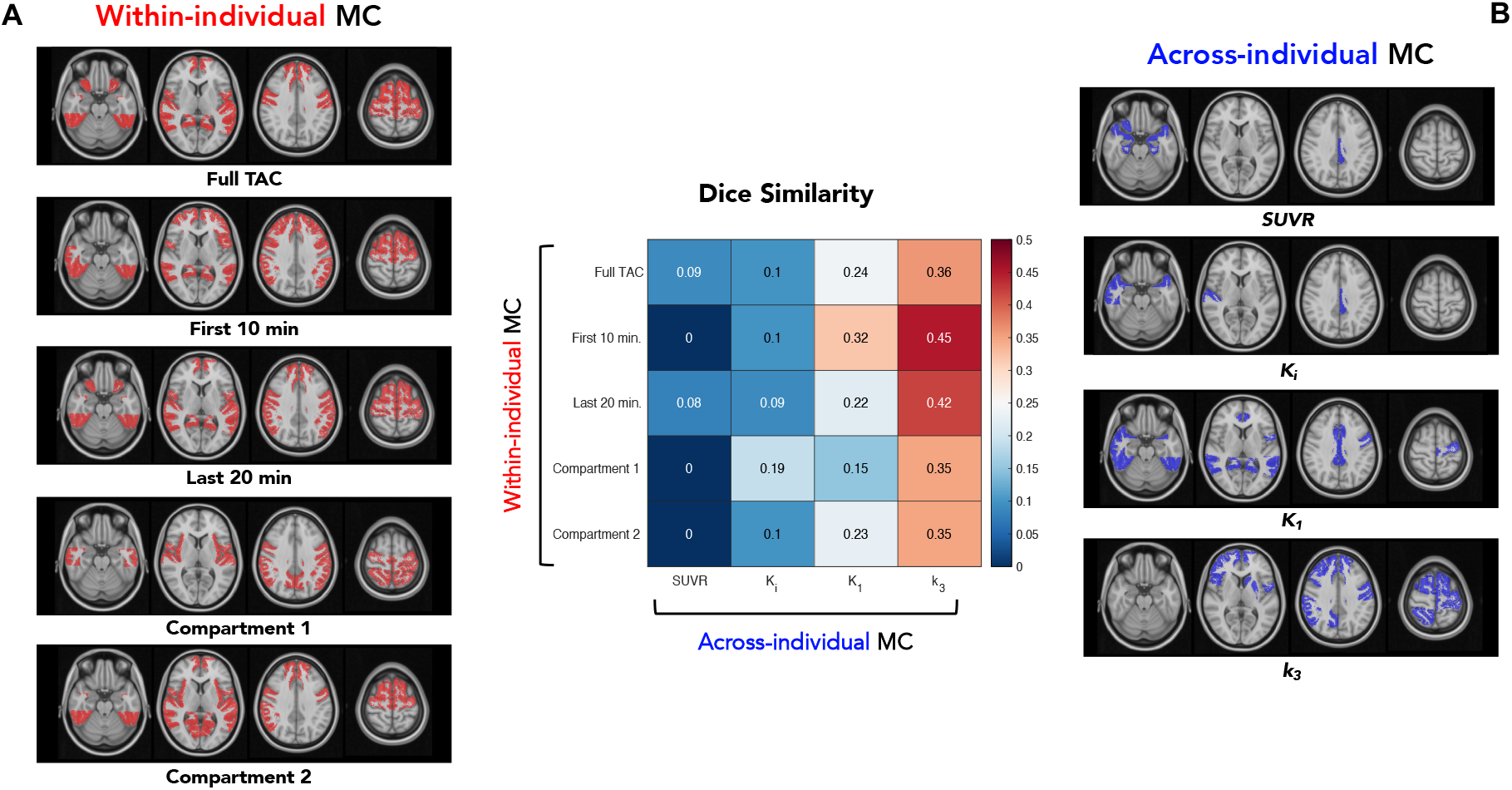
Comparison of wi-MC (**A**) vs. ai-MC (**B**) ‘hubs’, identified as the top DEG and EC nodes for each matrix. Hub nodes are shown on the Hammers atlas regions in red (**A**) and blue (**B**) respectively. A matrix of Dice similarity values between wi-MC and ai-MC hubs is shown in the central panel. Early-time and latetime dynamic PET MC have the highest association with across-subject covariation of *k_3_*.

As to ai-MC hubs, while *SUVR, K_i_* and *K_1_* have a similar hub distribution, mainly in temporal, insular and cingulate cortices, *k_3_* hubs fall in frontal and subcortical areas (**Figure 4B**).

When we look at the Dice similarity between hub vectors of wi-MC vs. ai-MC matrices, again we find a lack of match between *SUVR* MC and wi-MC hubs, with higher overlap in the case of *K_1_* and especially *k_3_*.

We also related the regional EC values from both wi-MC and ai-MC matrices to the average [^18^F]FDG *SUVR, K_i_, K_1_, k_3_* (**Supplementary Results**).

### Matching within-individual and across-individual MC with structural and functional connectivity

Finally, we assessed the similarity between the wi-MC and ai-MC matrices and a) a SC template (**Supplementary Figure 11A**), b) the group-average FC from the same individuals (**Supplementary Figure 11B**), to understand if a) structural or b) BOLD-based functional ‘connections’ could support the identified metabolic networks.

When looking at SC (**Figure 5A**), Dice similarity coefficients (DSC) tend to be higher for wi-MC matrices, especially for the *early* part of the TAC (DSC = 0.42), full TAC (DSC = 0.40) and *C*_1_ (*t*) (DSC = 0.40). Amongst the ai-MC matrices, *k_3_* has the highest similarity (DSC = 0.40), while *SUVR* the lowest (DSC = 0.31). Notably, the results are highly similar with another SC template (**Supplementary Figure 12**).

**Figure 5:**
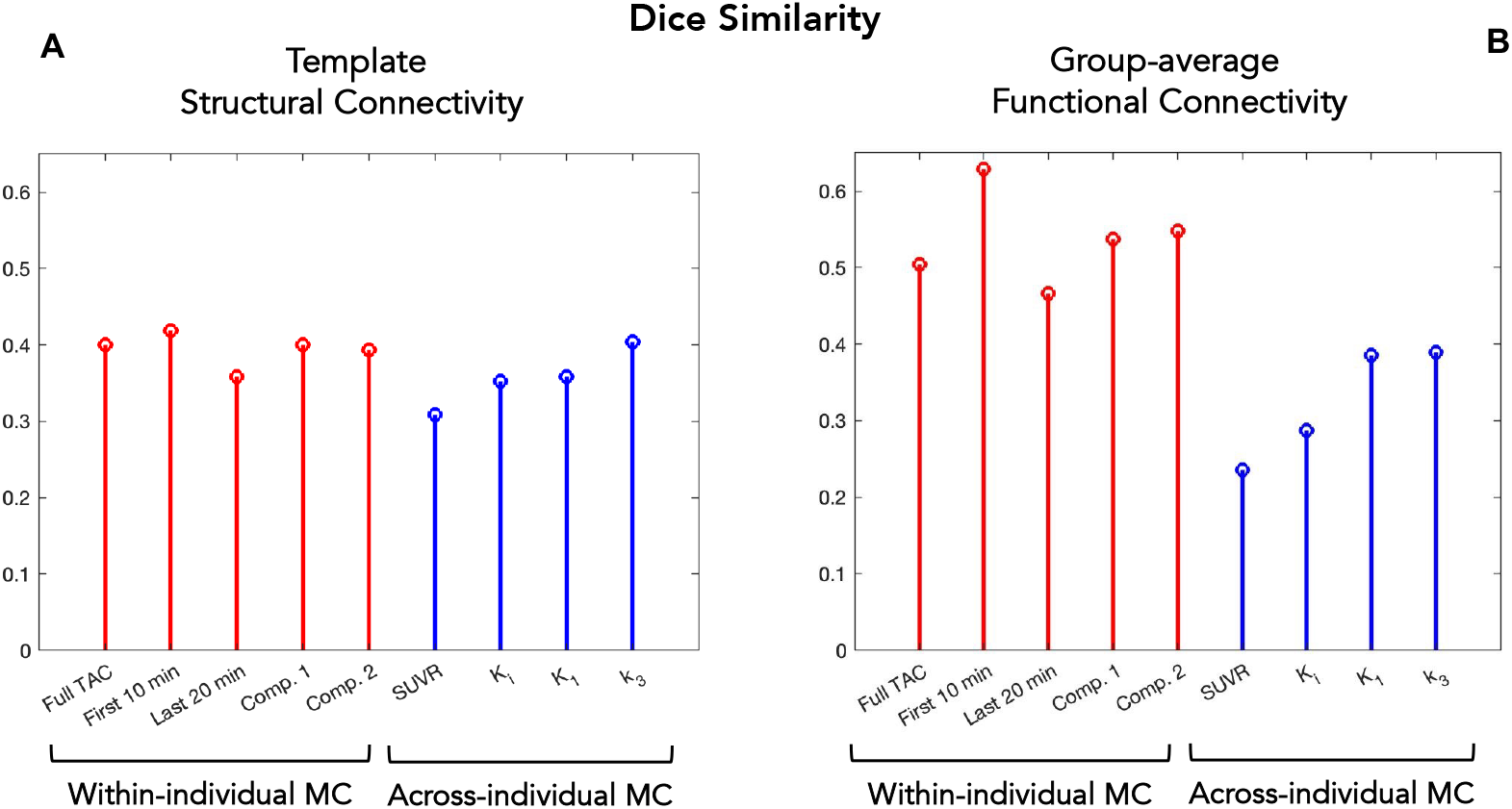
Dice similarity (stem plots) of the wi-MC (*red*) and ai-MC (*blue*) binarized matrices (80^th^ percentile) with the SC template (**A**) and group-average FC matrix (**B**).

When we look at the match with FC, the wi-MC matrices have even higher Dice (early TAC: DSC = 0.63, *C*_1_ (*t*): DSC = 0.54, *C*_2_(*t*): DSC = 0.55), while ai-MC maintain lower values (*k_3_*: DSC = 0.39, *SUVR:* DSC = 0.24) (**Figure 5B**). This result is reproduced also with the Schaefer atlas (full TAC: DSC = 0.44; early TAC: DSC = 0.38, *C*_1_(*t*): DSC = 0.42, *C*_2_(*t*): DSC = 0.40; *SUVR:* DSC = 0.35; *k_3_*: DSC = 0.33).

## Discussion

In this work, we reassessed the concept of metabolic connectivity from a PET kinetic modelling perspective, trying to capitalize on the multifaceted physiological information provided by dynamic [^18^F]FDG data.

The first issue we wanted to tackle was to select a good approach to estimate individual-level MC (i.e., wi-MC).

The methods used in the fMRI literature to assess individual-level FC conventionally rely on variance-based methods (correlation/covariance), designed to identify signals that vary together over time [3]. These approaches are also borrowed in the few studies which attempt to estimate individual-level MC from dynamic PET [6, 7, 9]. However, variance-based methods are problematic with dynamic PET data: while without any signal standardization no clear network structure is evident, the use of standardization produces different networks depending on the way the global signal is removed (**Supplementary Figure 2**). Importantly, signal fluctuations, which are at the core of resting-state fMRI measurements, are of unclear physiological meaning in the case of dynamic PET, even though in previous work [6, 7, 9] structured networks have been extracted from these fluctuations (reconstructing data with a uniform time grid which does not truly respect PET count statistics).

While it is remains unclear whether fluctuations contain relevant biological information or are more affected by non-physiological noise, the signal itself is for sure biologically informative, which is why it is used for kinetic modelling [2].

With respect to this, our method of choice (Euclidean similarity) identifies signals that are close in a Euclidean distance sense. When used on voxel-wise dynamic data, Euclidean distance already proved to effectively identify biologically meaningful clusters [29]. Here, we have repurposed this metric from a hard cluster assignment to a continuous space of *pharmacokinetic similarities* across brain regions, which, interestingly, display the hallmarks of brain connectivity studies, i.e., 1) a small-world network structure with 2) strong homotopy [3].

Moreover, the *between-individual variability* of the Euclidean similarity-based wi-MC matrices is low, which highlights the robustness of the chosen approach. This is the reason why we feel confident with looking at the *group-average* matrices, without risk of overlooking strong individual peculiarities as in fMRI FC [35].

We have proposed several types of MC, both for the within-individual and for the across-individual approaches.

To try and disentangle the multifaceted physiological information of dynamic PET data we have, for the first time, derived five different wi-MC matrices: the MC networks based on the early TAC and the time course of the free intracellular tracer concentration, *C*_1_(*t*), are expected to be more related to *inflow and delivery*, while the late TAC and the metabolized intracellular tracer concentration, *C*_2_(*t*), are more directly associated with *metabolic exchanges*. This overcomes the limitation of using only the raw tissue TAC, which combines the delivery and the actual metabolism [4]. We find good overlap between the late TAC and *C*_2_ (*t*), while the *C*_1_(*t*) wi-MC is the least similar to the other matrices, implying that kinetic modelling might provide non-redundant information for individual-level MC calculation.

Notably, this framework could easily be extended to other PET tracers, separating delivery, specific and non-specific binding to estimate individual-level connectivity for receptor systems [4].

We have also provided a new perspective on the dominant across-individual MC approach [4], always based on *SUVR*, due to it being the easiest parameter to obtain. In the case of other PET tracers, kinetic model parameters (e.g., *V_T_, BP_ND_*) have already been employed for ai-MC analyses [4, 36], but this is a new attempt for [^18^F]FDG. As we have shown, relevant differences emerge when the chosen parameter is not *SUVR*, especially with *k_3_*. *SUVR* and *K_i_* are typically considered to be the most important (and convenient) [^18^F]FDG parameters to summarize [^18^F]FDG metabolism, but this adds to the evidence that the microparameters (e.g., *K_1_* and *k_3_*) might bear important meaning, both in physiology [37] and pathology [38].

Our final aim was to enhance the interpretability of MC measures by relating the *within*- vs. *across-individual* MC frameworks to one another and to other connectivity measures, in a preliminary attempt at validation. With the *SUVR* ai-MC being the most employed approach, it is highly relevant to understand whether the *across-individual* estimates also reflect *individual-level* information.

When we assessed the match between the ai-MC and wi-MC, weak-to-moderate correlations are found for *K_1_*- and *k_3_*-based MC vs. wi-MC matrices, consistently with microparameters being more sensitive to the physiological processes probed by wi-MC, as both are a function of the same dynamic PET signal. The correlation with *SUVR* MC is instead low. Also, wi-MC hubs are more concentrated in frontotemporal areas, while ai-MC hubs of *SUVR, K_i_* and *K_1_* in temporal and limbic cortex. Notably, wi-MC hubs from infusion protocols were also in frontotemporal regions [6], unlike *SUVR* MC hubs [6][39].

Finally, we compared MC to typical measures of brain connectivity, i.e., SC and FC. There is not a high overlap between MC and SC (Dice similarity ~ 0.3-0.4). However, we found a higher match for wi-MC, and *SUVR* ai-MC (previously related to SC in [12]), is the one with the *least* similarity with the underlying structural network. Importantly, these results are consistent across different SC templates. To some extent, we can expect structural and functional/metabolic connections to have only a partial overlap, as already shown for SC vs. FC [15], since structural connectivity highlights only *direct* connections (plus technical issues which may cause SC estimates to miss some tracts). A more fine-grained investigation, focusing on short- vs. long-range or within- vs. between-network connections, might bear additional insight [12].

The match between FC and wi-MC is good (Dice ~ 0.5) with both atlases, and higher than the FC vs. ai-MC match and the wi-MC vs. ai-MC match itself (**Table 1**). This is in line with bolus and infusion findings [6, 9]. While further investigation of the FC-MC coupling at a finer scale is required, our findings show how individual-level MC captures more of the BOLD-based functional network information than its across-individual counterpart. It is intriguing to wonder which reasons might be behind the correspondence between *pharmacokinetic similarity* among brain regions and FC, but it seems the metabolic properties underlying “hubness” in the MC network might inherently be linked to a region’s capacity to connect with other regions, both locally and globally. This also indicates that the coupling between glucose metabolism and fMRI FC, which we found to be somewhat limited when considering only *local* metabolic measures like *SUVR* [40], may become stronger when *both* PET and rs-fMRI are framed into a *large-scale* connectivity environment.

In conclusion, can within-individual MC be seen as a preferable approach? Overall, our comparison between wi-MC and ai-MC shows that individual-level estimates provide non-redundant networks which cannot be inferred from ai-MC, in agreement with what was found for ai-MC vs. *infusion* wi-MC [6]. Moreover wi-MC seems to be more interpretable in terms of match with fMRI FC. Together, these properties suggest a potential for individual PET connectivity measures to be used as a biomarker, which warrants further exploration.

There are limitations to this work. Absolute quantification of Sokoloff’s model parameters requires an input function ideally obtained via arterial sampling [2]. Bias in the estimates might have been introduced by the IDIF approach, especially since the data come from a low-spatial-resolution HR+ scanner, but we believe our multi-step IDIF extraction algorithm with Chen’s correction [18] (the reference standard in IDIF extraction with older scanners, and the only approach with potential for calculating microparameters [41]) allows to retrieve sufficiently accurate inputs. Moreover, for MC estimates, it is not the *absolute* parameter value that is of interest, but their *relative* spatial distribution, which is likely to be preserved. Also, our Bayesian estimation framework minimizes noise [26]. Future work should reassess these findings using PET data with higher spatiotemporal resolution [38].

Our approach does not solve the inherent problem of functional network analyses: connectomes may have plausible structure, but their biological underpinnings and interpretation remain more elusive since a “ground truth” is rarely available. The attempts at validation presented here (relating MC to other connectivity measures) provide only a partial understanding of the underlying mechanisms. Further efforts are highly warranted, possibly using interventional approaches in animal models to elucidate causative links.

## Conclusion

After developing a new approach to estimate individual-level wi-MC using considerations of tracer kinetics, and estimating across-individual ai-MC from [^18^F]FDG kinetic parameters, we thoroughly assessed the relationships between wi-MC and ai-MC at multiple levels, i.e., in terms of matrix and hub similarity, and their match with FC and SC matrices. We found the two MC frameworks to provide different and somewhat complementary information, with wi-MC having higher similarity with the FC network. Future work should consider applying wi-MC approaches to clinical populations, to verify if they can provide useful biomarkers.

## Statements & Declarations

### Author Contributions

Andrei G. Vlassenko and Tony Durbin collected the data. Tommaso Volpi and Alessandra Bertoldo designed the research. Tommaso Volpi, Giulia Vallini, Erica Silvestri, and Mattia De Francisci analyzed the data. Tommaso Volpi, Erica Silvestri, Maurizio Corbetta, John J. Lee, Andrei G. Vlassenko, Manu S. Goyal and Alessandra Bertoldo interpreted the results. Tommaso Volpi wrote the manuscript. All authors revised the manuscript.

### Consent to participate

All assessments and imaging procedures were approved by Human Research Protection Office and Radioactive Drug Research Committee at Washington University in St. Louis. Written consent was provided from each participant.

