## Supplementary Materials for "A new framework for metabolic connectivity mapping using bolus [^18^F]FDG PET and kinetic modelling"

*European Journal of Nuclear Medicine and Molecular Imaging*

### Supplementary Methods

*Data acquisition*

For each participant, a structural MRI scan was performed to provide anatomical information. High-resolution structural images were acquired on a Siemens Magnetom Prisma$^{fit}$ scanner using a 3D sagittal T1-weighted magnetization-prepared 180° radio-frequency pulses and rapid gradient-echo (MPRAGE) multi-echo sequence (TE = 1.81, 3.6 , 5.39, 7.18 ms, TR = 2,500 ms, TI = 1,000 ms, 0.8×0.8×0.8-mm voxels). The final T1w image was obtained as the average of the first two echoes [1]. Additionally, T2* gradient-echo echo planar imaging (GE-EPI) data were acquired (TR/TE=800/33 ms, flip angle 52°, voxel size 2.4×2.4×2.4 mm, MB 6, 375 volumes for total scan time of 5 min), together with two spin-echo (SE) acquisitions (TR/TE=6000/60 ms, flip angle 90°) with opposite phase encoding directions (AP, PA).

[$^{18}$F]FDG scans were performed on a Siemens model 962 ECAT EXACT HR + PET scanner (Siemens/CTI) [2], as previously described [3], after i.v. bolus injection of 5.1 ± 0.3 mCi (187.7 ± 12.1 MBq) of [$^{18}$F]FDG. Dynamic acquisition of PET emission data continued for 60 min.

Participant head movements during scanning were restricted by a thermoplastic mask. All PET images were acquired in the eyes-closed waking state. No specific instructions were given regarding cognitive activity during scanning other than to remain awake.

PET data were reconstructed via filtered back-projection (ramp filter, 5 mm FWHM) as 128x128x63 matrices. Attenuation correction was performed using the participant's transmission scan. The chosen reconstruction grid consisted of 52 frames of increasing duration (24 x 5 s frames, 9 x 20 s frames, 10 x 1 min frames, and 9 x 5 min frames).

Venous samples for plasma glucose determination were obtained just before and at the midpoint of the scan to verify that glucose levels were within normal range throughout the study. Also, venous samples were collected to assess [$^{18}$F]FDG plasma concentration, with two possible sampling schedules: for most participants, sampling occurred 20, 30, 45 minutes after injection of the radiotracer, whereas, for a minority of participants (n = 10), samples were acquired after 30, 40 and 50 minutes. Each sample consisted of about 2 ml, half of which was used to measure radioactivity in plasma. Radioactivity counter measurements was given in counts per 12 seconds. The counter's efficiency (0.2707 cps/Bq) was experimentally determined [4].

*Structural MRI preprocessing*

Structural T1w images were N4 bias field-corrected [5], skull-stripped [6], and segmented into grey matter (GM), white matter (WM) and cerebrospinal fluid (CSF) using SPM12 [7].

T1w images were normalized to the symmetric MNI152 2009c atlas [8] via nonlinear diffeomorphic registration [9].

The Hammers anatomical atlas [10] and the Schaefer functional atlas (100 ROIs, 7 networks) [11] were registered to T1w space by inverting the obtained nonlinear transformation.

*Functional MRI preprocessing*

The fMRI data were analyzed in a similar way to the Human Connectome Project minimal preprocessing pipeline [12].

The first four fMRI volumes were discarded to avoid non-equilibrium magnetization effects. The remaining volumes were corrected for slice timing differences [13] and magnetic field distortion [14], and realigned to the median volume [15]. A template EPI volume [16] was obtained from realigned fMRI data and used to estimate an affine transform employed to map main tissue segmentations obtained from the T1w image to the EPI space.

Nuisance signals, including motion parameters and their first order derivatives, and the first 5 temporal principal components obtained after principal component analysis of WM and CSF EPI signals as in [17], were regressed out from all brain voxels in native EPI space. Finally, the BOLD signal was high-pass filtered with a cut-off of 0.008 Hz.

*PET kinetic modelling*

A static PET image was obtained by summing late PET frames (40-60 min) after motion correction. The static PET image was normalized into *SUVR* by dividing each voxel's value by the whole-brain [$^{18}$F]FDG average uptake [18].

To perform PET kinetic modelling, an image-derived input function (IDIF) was extracted from dynamic PET data using a semi-automatic pipeline [19]:

- segmentation of the internal carotid arteries was performed on a pseudo-angiography image (obtained by summing dynamic PET frames up to an adaptive threshold of one frame before the peak time for venous vessels), on which a *vesselness* algorithm (Jerman filter) [20] was run to generate a vessel mask;
- selection of "hot voxels" within the mask, according to their peak amplitude and time-to-peak;
- parametric clustering [21] (k-means algorithm, k = 2, squared Euclidean distance, 500 replicates) on seven parameters calculated on the TAC of each voxel (peak amplitude, slope of rising part before peak, slope of tail, area under the curve before and after the peak, tail average value, TAC standard deviation), with the cluster having the highest peak centroid being selected and used to derive the raw IDIF;
- IDIF model fitting was performed using a modified version of Feng's model [22, 23] with maximum a posteriori estimation of the exponential decay parameters;
- Chen's spillover correction [24] was applied to the fitted IDIF curves using three venous samples (obtained after arteriovenous equilibration, i.e., after 20 min post-injection) and a background tissue TAC, obtained as the highest activity cluster centroid within a background mask (obtained from morphological dilation of the *vesselness* mask);
- IDIF shift correction, to correct for delay between the carotids and the voxel of interest.

Voxel-wise estimation of Sokoloff's model parameters was performed using a Variational Bayesian approach [25], according to the following pipeline:

- a k-means clustering approach was applied to the dynamic PET data, extracting 6 GM and 5 WM clusters (as from the tissue segmentations linearly mapped to PET space),
- conventional nonlinear estimation of Sokoloff's model (using weighted nonlinear least squares, with weights chosen as the inverse of the variance of the PET measurement error [26]) was performed at the regional level, i.e., on the 11 cluster centroids,

- voxel-wise estimation of the model parameters via Variational Bayesian inference was performed using prior distributions derived from cluster-wise estimates.

We refer to [25] for more complete details on the parameter estimation procedure.
Parametric maps (i.e., voxel-wise maps) of $K_1$ [ml/cm$^3$/min] (inflow of tracer), $k_2$ [min$^{-1}$] (efflux of tracer), $k_3$ [min$^{-1}$] (phosphorylation of tracer), $V_b$ [%] (blood volume fraction) were obtained for each participant. The parametric map of $K_i$ [ml/cm$^3$/min] (irreversible uptake of tracer) was obtained by the solving the following equation at voxel level:

$$K_i = \frac{K_1 k_3}{k_2 + k_3} \qquad (1)$$

The group-average maps of *SUVR*, $K_i$, $K_1$, $k_3$ are shown in **Supplementary Figure 6**.
Importantly, we did not perform partial volume correction on PET data: the use of the GM segmentation provided by SPM, plus avoiding spatial smoothing of PET data during processing allowed to minimize PVEs, as also suggested in recent PET analysis works [27, 28, 29]. Moreover, there is no gold standard for partial volume correction, especially in dynamic PET studies, as this procedure is known to affect the accuracy of kinetic modeling and potentially alter spatial metabolic patterns [30, 31].
The voxel-wise predictions of the time-varying free intracellular concentration of the [$^{18}$F]FDG tracer ($C_1(t)$, [kBq/cm$^3$]) and its metabolized (i.e., [$^{18}$F]FDG 6-phosphate) intracellular concentration ($C_2(t)$, [kBq/cm$^3$]), representative of hexokinase activity, were reconstructed from Laplace transform solutions of Sokoloff's model [**Error! Reference source not found.**] b y using the IDIF as plasma input function ($C_p(t)$ [kBq/ml]):

$$C_1(t) = \frac{K_1 k_2}{k_2 + k_3}\, e^{-(k_2+k_3)t} \otimes C_p(t) \qquad (2)$$

$$C_2(t) = K_i \int_0^t C_p(\tau) d\tau \qquad (3)$$

*Within-individual metabolic connectivity (wi-MC)*

The final approach presented in the paper is based on the Euclidean distance $d_{x_1,x_2}$ between each pair of TACs $x_{i,1}$ and $x_{i,2}$:

$$d_{x_1,x_2} = \sqrt{\sum_{i=1}^{T} \left(x_{i,1} - x_{i,2}\right)^2} \text{ with } T = \text{number of time points} \qquad (4)$$

Notably, distance-based approaches are frequently employed to perform cluster analysis on dynamic PET data [32]. From $d_{x_1,x_2}$we derived a measure of Euclidean similarity, as 1 minus the normalized $d_{x_1,x_2}$, i.e., divided by the maximum distance among pairs of TACs, and thus scaled to [0, 1]. Due to the markedly heavy-tailed (left-skewed) distribution of Euclidean similarity ES values, a Fisher z-transformation was applied, and then the values were again rescaled to [0, 1].

*List of the Hammers ROIs used in the main analyses:*

**LEFT HEMISPHERE**
**Subcortical**
1. Cerebellum
2. Caudate nucleus
3. Nucleus accumbens
4. Putamen
5. Thalamus
6. Pallidum

**Frontal**
7. Middle frontal gyrus
8. Precentral gyrus
9. Straight gyrus
10. Anterior orbital gyrus
11. Inferior frontal gyrus
12. Superior frontal gyrus
13. Medial orbital gyrus
14. Lateral orbital gyrus
15. Posterior orbital gyrus
16. Subgenual frontal cortex
17. Subcallosal area
18. Pre-subgenual frontal cortex

**Insula & Cingulum**
19. Insula
20. Cingulate gyrus (gyrus cinguli), anterior part
21. Cingulate gyurs (gyrus cinguli), posterior part

**Temporal**
22. Hippocampus
23. Amygdala
24. Anterior temporal lobe, medial part
25. Anterior temporal lobe, lateral part
26. Parahippocampal and ambient gyri
27. Superior temporal gyrus, posterior part
28. Superior temporal gyrus, posterior part
29. Fusiform gyrus
30. Posterior temporal lobe
31. Superior temporal gyrus, anterior part

**Parietal**
32. Postcentral gyrus
33. Superior parietal gyrus
34. Inferiolateral remainder of parietal lobe

**Occipital**
35. Lingual gyrus
36. Cuneus
37. Lateral remainder of occipital lobe

**RIGHT HEMISPHERE**
**Subcortical**
38. Cerebellum
39. Caudate nucleus
40. Nucleus accumbens
41. Putamen
42. Thalamus

43. Pallidum

**Frontal**

44. Middle frontal gyrus
45. Precentral gyrus
46. Straight gyrus
47. Anterior orbital gyrus
48. Inferior frontal gyrus
49. Superior frontal gyrus
50. Medial orbital gyrus
51. Lateral orbital gyrus
52. Posterior orbital gyrus
53. Subgenual frontal cortex
54. Subcallosal area
55. Pre-subgenual frontal cortex

**Insula & Cingulum**

56. Insula
57. Cingulate gyrus (gyrus cinguli), anterior part
58. Cingulate gyurs (gyrus cinguli), posterior part

**Temporal**

59. Hippocampus
60. Amygdala
61. Anterior temporal lobe, medial part
62. Anterior temporal lobe, lateral part
63. Parahippocampal and ambient gyri
64. Superior temporal gyrus, posterior part
65. Superior temporal gyrus, posterior part
66. Fusiform gyrus
67. Posterior temporal lobe
68. Superior temporal gyrus, anterior part

**Parietal**

69. Postcentral gyrus
70. Superior parietal gyrus
71. Inferiolateral remainder of parietal lobe

**Occipital**

72. Lingual gyrus
73. Cuneus
74. Lateral remainder of occipital lobe

**Supplementary Results**

After relating the wi-MC and ai-MC approaches to one another, we tried to assess the level of similarity between information derived from these networks and more physiologically interpretable parameters, i.e., those derived from [$^{18}$F]FDG PET quantification.
In particular, the eigenvector centrality (EC) graph metric, which describes the level of 'connectedness' of a region in each MC network, was calculated on both wi-MC and ai-MC matrices, and the EC values were plotted against the across-individual mean values of [$^{18}$F]FDG $SUVR$, $K_i$, $K_1$, $k_3$ (**Supplementary Figure 10**).

While the EC of ai-MC matrices (**Supplementary Figure 10B**) have overall weak relationships with [$^{18}$F]FDG parameters, typically with a negative sign, the EC of wi-MC matrices (**Supplementary Figure 10A**) have positive relationships with the parameters, which tend to be highly nonlinear (well related to the wi-MC through a quadratic relationship) especially for the *full* TAC, *late* part, and $C_2(t)$ vs. $SUVR$.
This relationship is preserved on the Schaefer atlas.

### Supplementary Discussion

*Within-individual metabolic connectivity*

Of note, little overall variability in wi-MC matrices is demonstrated if we take age into account and compare the matrix obtained from the older vs. the younger individuals (r values between 0.86 and 0.93, $p < 10^{-9}$, Bonferroni corrected).

Further work is necessary to understand the biologically meaningful sources of within- and between-individual variability in our wi-MC estimates.

*Comparing within-individual vs. across-individual MC: similarity of matrices and hubs, match with [$^{18}$F]FDG parameters*

When relating average [$^{18}$F]FDG parameters to the MC eigenvector centrality (a measure of global 'connectedness' which was found to summarize multi-scale neural properties [33]) across regions (**Supplementary Figure 10**), we find a strong nonlinear and non-monotonic relationship for wi-MC, while for ai-MC absent or negative relationships are detected, which are harder to interpret.

*Matching within-individual and across-individual MC with structural and functional connectivity*

We have noted that *subcortical areas* in our wi-MC matrices display lower 'connectivity' than the cortical ones. This seems to parallel the lower BOLD FC of subcortical regions. A direct comparison cannot be drawn, since subcortical BOLD signals are frequently sampled poorly, due to the need to optimize MR sensitivity to cortical areas [34], and our distance-based MC approach is different from the correlation-based fMRI FC, however, our results imply that subcortical PET kinetics are more dissimilar to one another than the cortical kinetics are, despite still highlighting relevant within-subcortex and cortico-subcortical relationships (**Supplementary Figure 4**), which warrant a more detailed assessment.

### Supplementary Figures

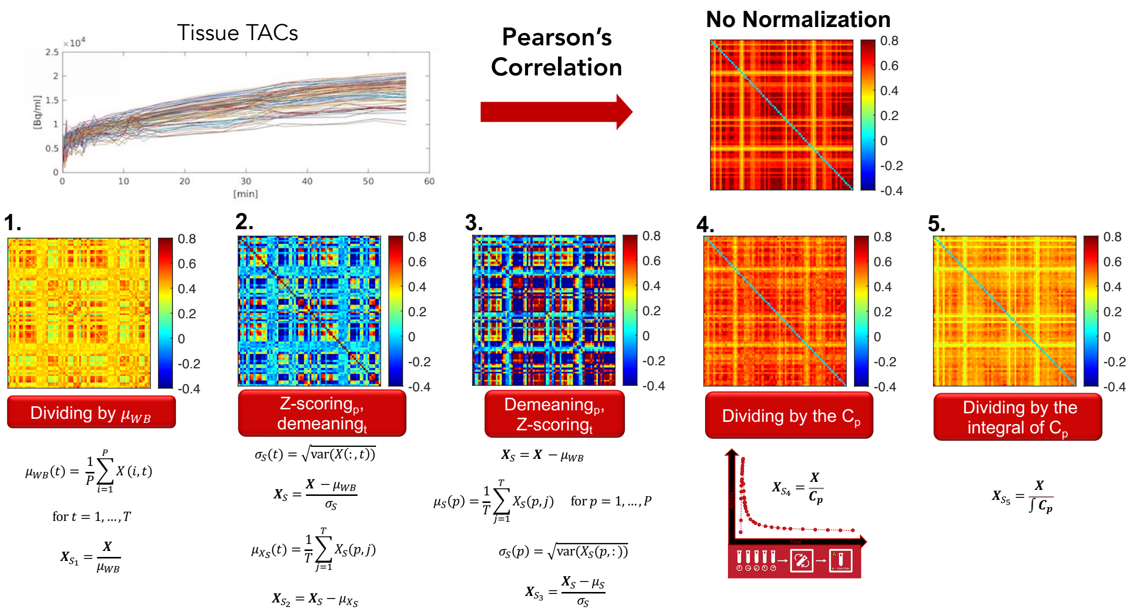


**Supplementary Figure 1**: Group-average *within-individual* MC matrices (Hammers atlas) obtained from the full tissue TAC, using Pearson's correlation as a similarity metric. The non-normalized case is compared with five different normalizations on the matrix of full tissue TACs $\boldsymbol{X} \in \mathbb{R}^{p \times T}$ (with feature size $p$ and sample size $T$): division by mean TAC $\mu_{WB}$ (1), z-scoring across regions followed by demeaning across time points (2), demeaning across regions (removing $\mu_{WB}$) followed by z-scoring across time (3), division by IDIF ($C_p$) curve (4), division by IDIF integral curve (5).

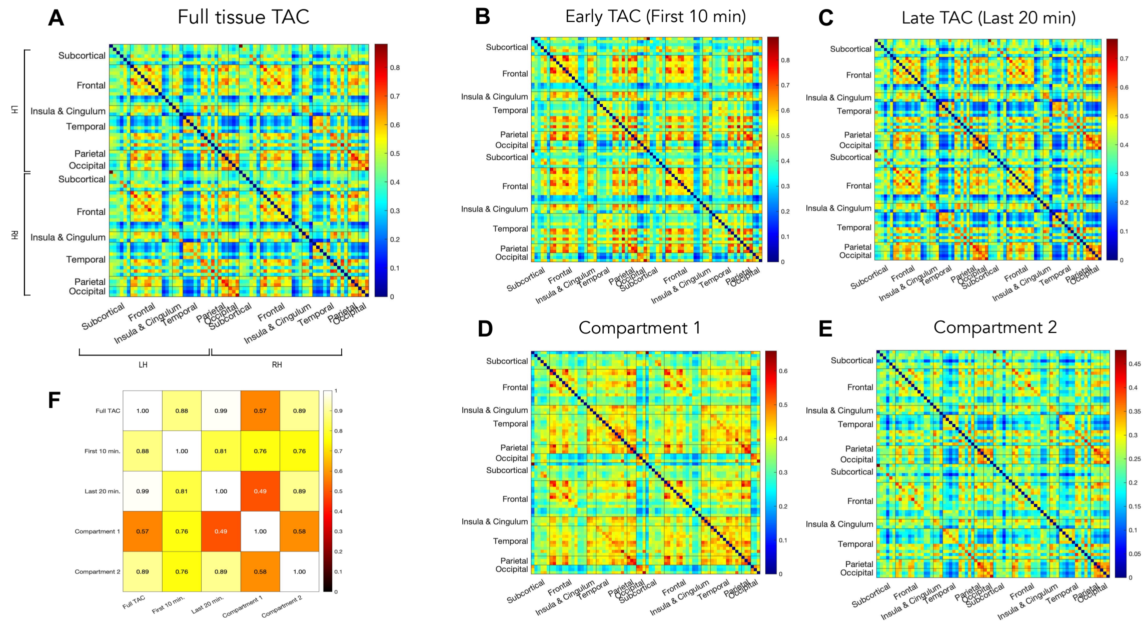


**Supplementary Figure 2**: *Within-individual* MC matrices (Hammers atlas). We report the group-average (n = 54) MC matrices obtained at the individual level from the *full* tissue TAC (**A**), its *early* part (**B**) and *late* part (**C**), the kinetics of $C_1(t)$ (**D**) and $C_2(t)$ (**E**), via the Euclidean similarity metric. We also report the Pearson's correlation matrix between the edges of the 5 wi-MC matrices (upper triangle) (**F**).

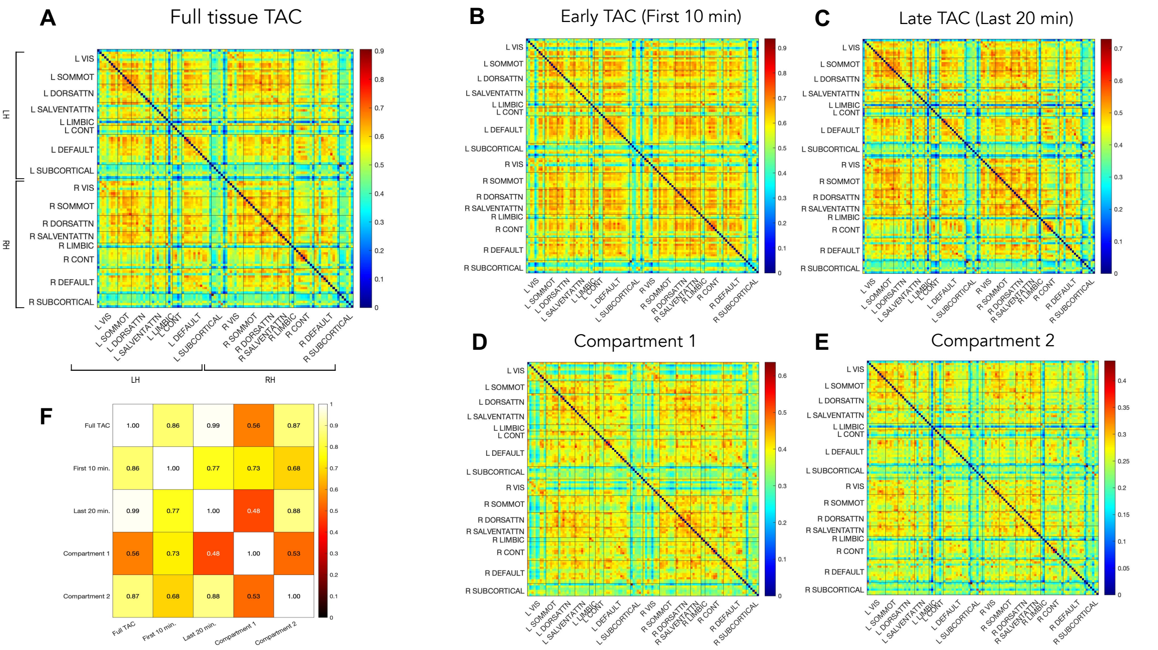


**Supplementary Figure 3**: *Within-individual* MC matrices (Schaefer atlas). We report the group-average (n = 54) MC matrices obtained at the individual level from the *full* tissue TAC (**A**), its *early* part (**B**) and *late* part (**C**), the kinetics of $C_1(t)$ (**D**) and $C_2(t)$ (**E**), via the Euclidean similarity metric. We also report the Pearson's correlation matrix between the edges of the 5 wi-MC matrices (upper triangle) (**F**).

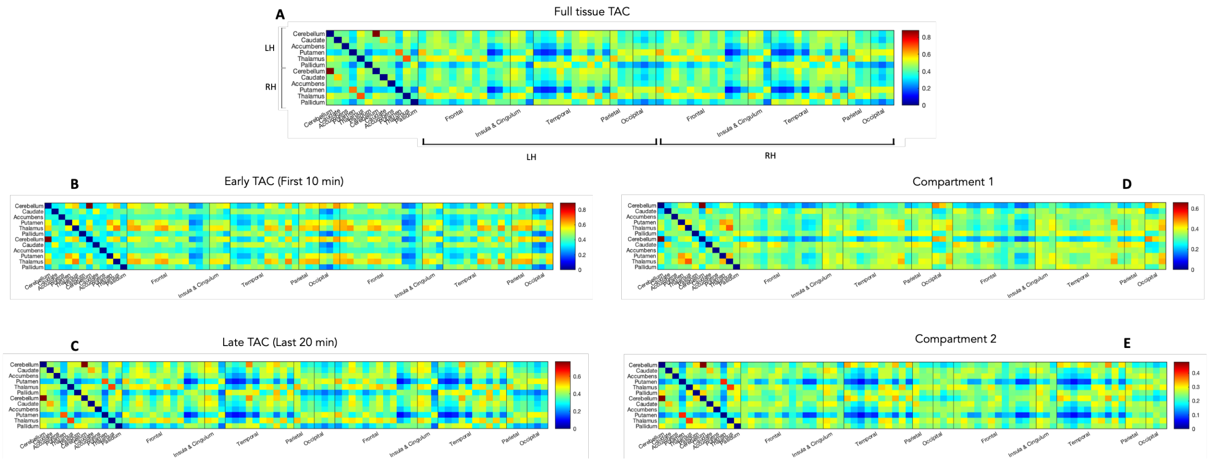


**Supplementary Figure 4**: *Within-individual* MC matrices – subcortical connectivity (Hammers atlas). We highlight the subcortical portion of the MC matrices obtained at the individual level from the *full* tissue TAC (**A**), its *early* part (**B**) and *late* part (**C**), the kinetics of $C_1(t)$ (**D**) and $C_2(t)$ (**E**), via the Euclidean similarity metric. From here, some relevant within-subcortex and cortico-subcortical relationships emerge, such as the moderate-strong coupling of the cerebellum with putamen, thalamus and especially the occipital cortex for $C_1(t)$ wi-MC, a finding which is reflected in the high $K_1$ values of these regions (**Supplementary Figure 6**).

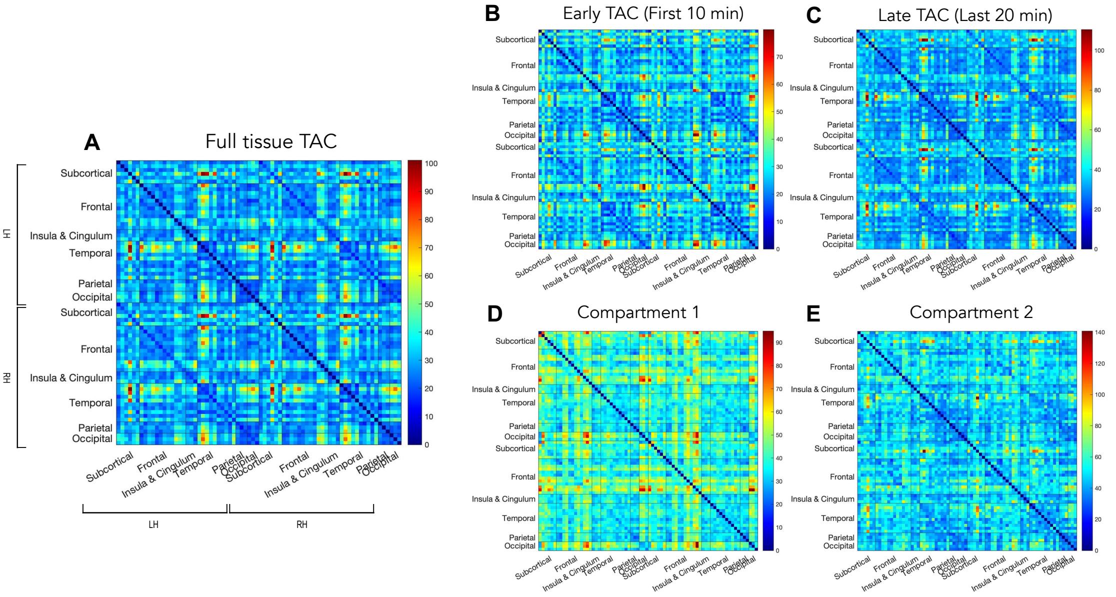


**Supplementary Figure 5**: Between-individual variability of *within-individual* MC matrices (Hammers atlas). We report the edge-level across-individual coefficients of variation (%) of MC matrices obtained from the *full* tissue TAC (**A**), its *early* part (**B**) and *late* part (**C**), the kinetics of $C_1(t)$ (**D**) and $C_2(t)$ (**E**).

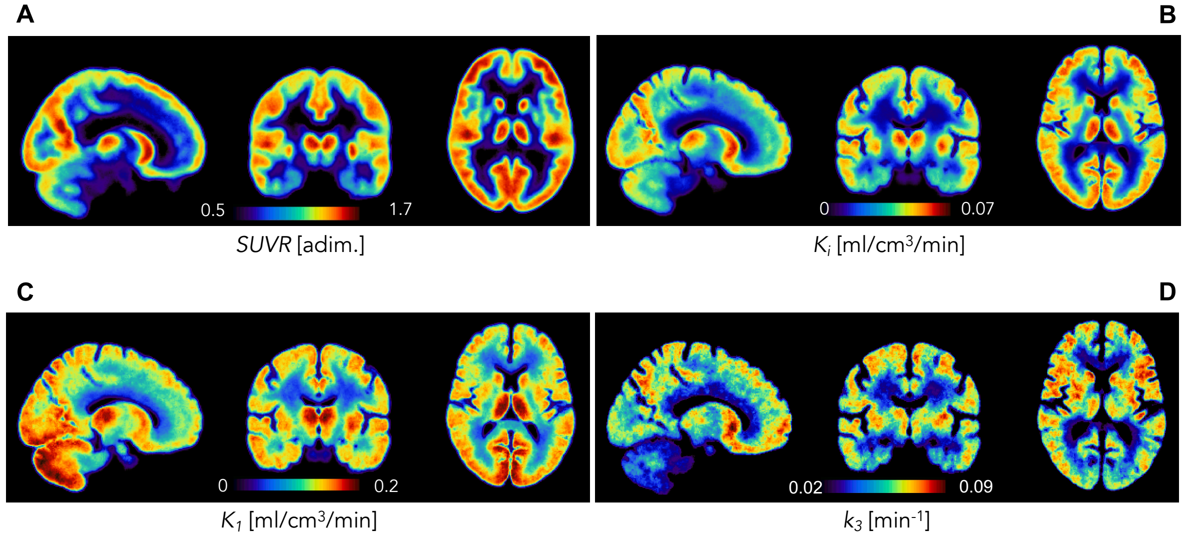


**Supplementary Figure 6**: Group-average [$^{18}$F]FDG parametric maps (n = 54) for *SUVR* (**A**), $K_i$ (**B**), $K_1$ (**C**) and $k_3$ (**D**) in MNI space.

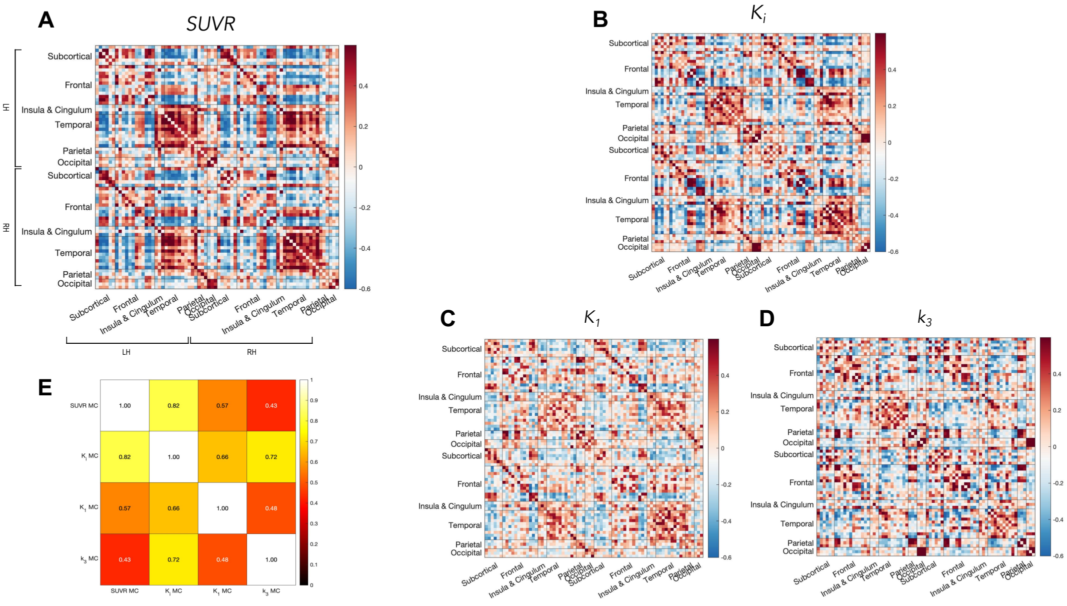


**Supplementary Figure 7**: *Across-individual* MC matrices (Hammers atlas). We report the across-individual Pearson's correlation matrices for *SUVR* (**A**), $K_i$ (**B**), $K_1$ (**C**) and $k_3$ (**D**). We also report the Pearson's correlation matrix between the edges of the 4 ai-MC matrices (upper triangle) (**E**).

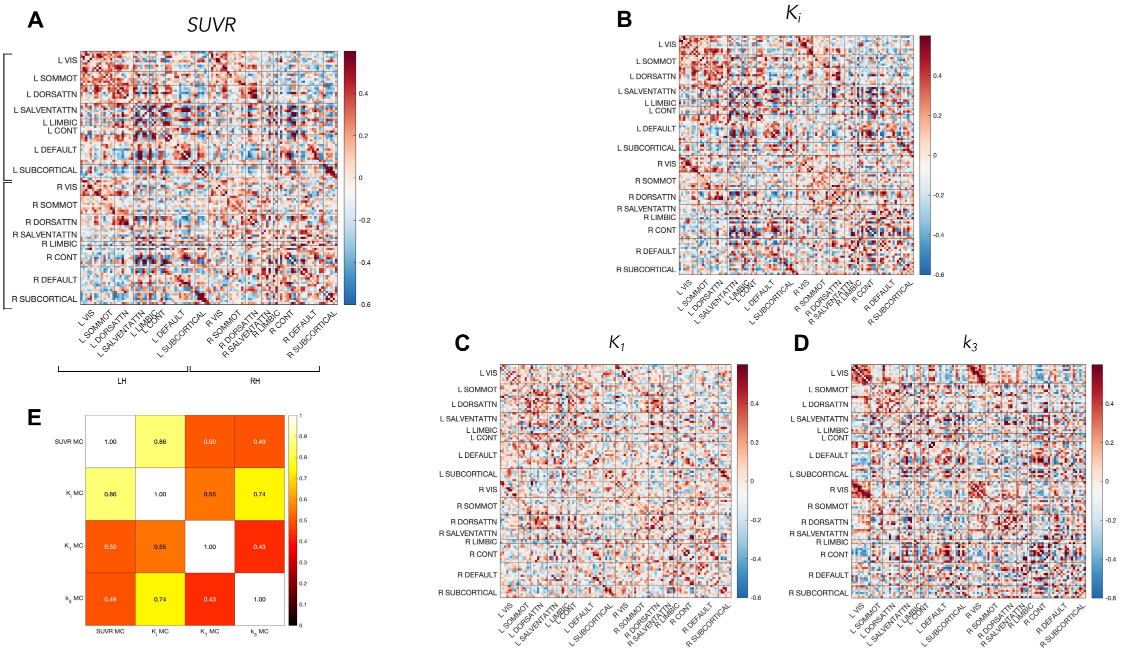


**Supplementary Figure 8**: *Across-individual* MC matrices (Schaefer atlas). We report the across-individual Pearson's correlation matrices for *SUVR* (**A**), $K_i$ (**B**), $K_1$ (**C**) and $k_3$ (**D**). We also report the Pearson's correlation matrix between the edges of the 4 ai-MC matrices (upper triangle) (**E**).

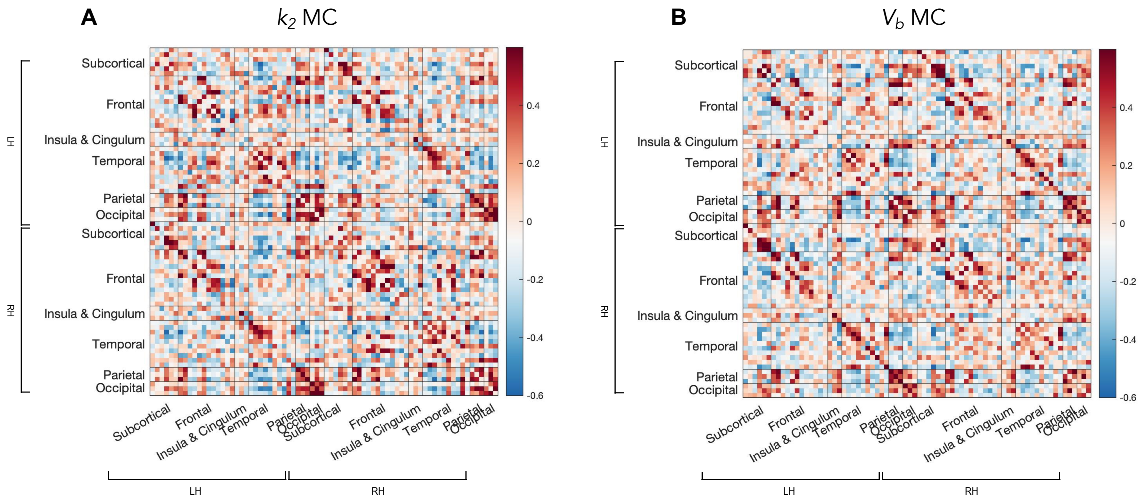


**Supplementary Figure 9**: *Across-individual* MC matrices for the two additional parameters obtained via compartmental modelling, i.e., $k_2$ [min$^{-1}$] (**A**), and $V_b$ [%] (**B**). Notably, they are both structured matrices, with an evident main diagonal and secondary diagonals.

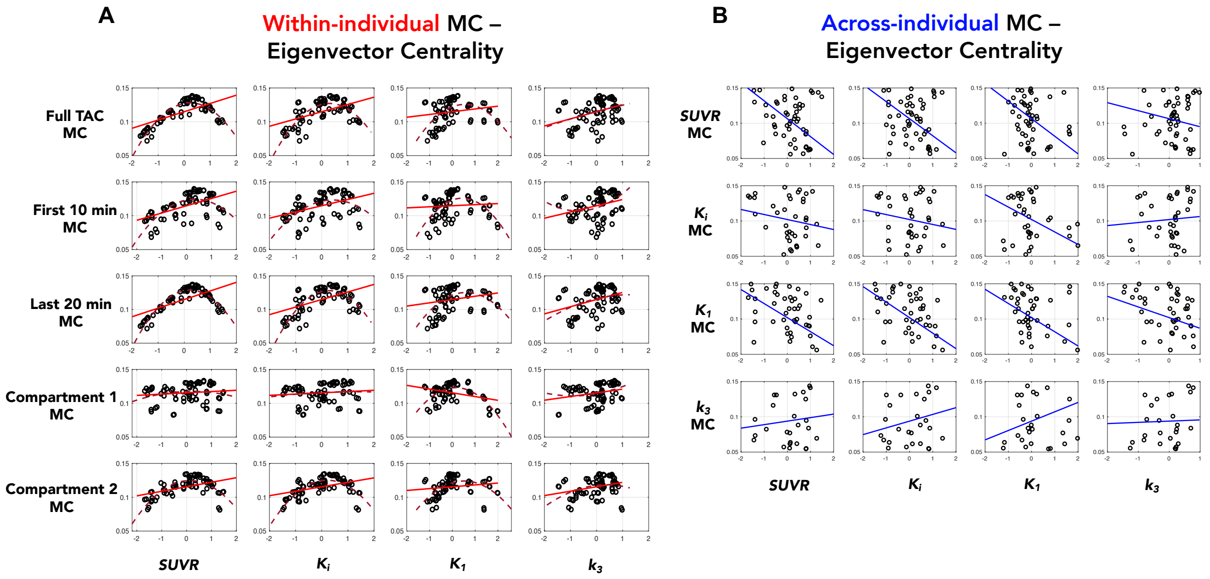


**Supplementary Figure 10**: Scatter plots of the across-region associations between group-average values of *SUVR*, $K_i$, $K_1$ and $k_3$ (z-scores, on the x axis) and the EC of *within-individual* (**A**) and *across-individual* MC (**B**) matrices (on the y axis). A linear fit line is shown in both **A** (red) and **B** (blue); a quadratic fit is shown as a red dashed line in **A**. A nonlinear and non-monotonic relationship is present for wi-MC, with regions that have high kinetic similarity to the rest of the brain (high EC) tending to have average glucose metabolism and delivery, and nodes whose TACs that are very dissimilar from the rest (low EC) having extreme metabolic rates (i.e., very high or very low); for ai-MC, instead, absent or negative relationships are detected, except for $k_3$ EC.

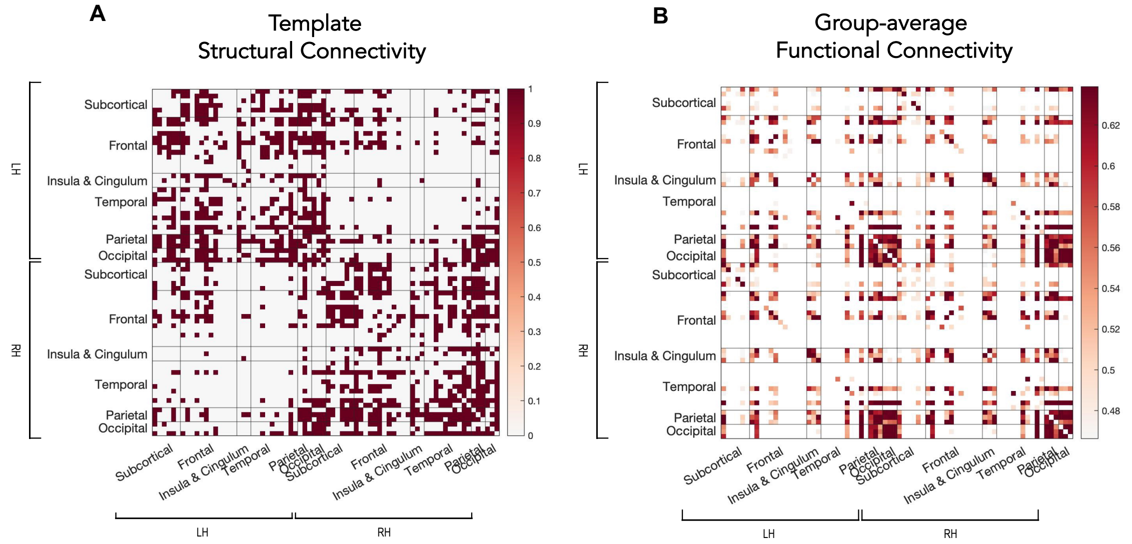


**Supplementary Figure 11**: Population-level connectomes employed for comparison with wi-MC and ai-MC. The binarized SC matrix (**A**) is obtained from averaging 178 structural connectomes. The group-average FC matrix (**B**), thresholded at the 80$^{th}$ percentile, is obtained from the same healthy individuals as the MC matrices (n = 54).

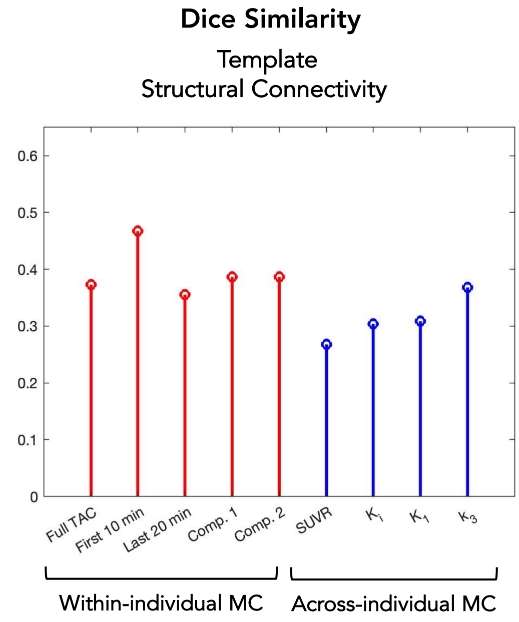


**Supplementary Figure 12**: Stem plot of the Dice similarity values between the group-average wi-MC (*red*) and ai-MC (*blue*) binarized matrices (80$^{th}$ percentile) and a second SC template.
